# CellMentor: Cell-Type Aware Dimensionality Reduction for Single-cell RNA-Sequencing Data

**DOI:** 10.1101/2025.06.17.660094

**Authors:** Or Hevdeli, Ekaterina Petrenko, Dvir Aran

## Abstract

Single-cell RNA sequencing (scRNA-seq) enables high-resolution profiling of individual cells, yet transforming this high-dimensional data into biologically meaningful representations remains a critical challenge. Current dimensionality reduction methods often fail to effectively balance technical noise reduction with preservation of cell-type-specific biological signals, particularly when integrating data across multiple experiments. Here, we present CellMentor, a novel supervised non-negative matrix factorization (NMF) framework that leverages labeled reference datasets to learn biologically meaningful latent spaces that can be transferred across related datasets. CellMentor employs a loss function that preserves cell type identity by simultaneously minimizing variation within known cell populations while maximizing distinctions between different cell types. We evaluated CellMentor against state-of-the-art dimensionality reduction and integration methods using controlled simulations of increasing difficulty and diverse real tissue types, each consisting of a labeled reference dataset and an unlabeled query dataset. In simulations, CellMentor maintained near-perfect clustering performance even under challenging conditions where other methods failed. In real datasets from PBMC, pancreas, and melanoma tissues, CellMentor demonstrated superior cell type separation while effectively mitigating batch effects. CellMentor also excelled at detecting rare cell populations and maintained reasonable performance when encountering novel cell types absent from reference data. With its robust batch correction capabilities and ability to preserve biologically meaningful cell type distinctions, CellMentor is particularly valuable for integrative analyses across multiple experiments.

## 1 INTRODUCTION

Single-cell RNA sequencing (scRNA-seq) is a powerful tool for investigating cellular heterogeneity, enabling analysis of gene expression at the single-cell level. This technology has greatly enriched our understanding of biological phenomena and disease mechanisms, revealing the diversity and function of cellular systems. The established scRNA-seq analytical workflow includes quality control and filtering, data normalization, feature selection, dimensionality reduction, and unsupervised clustering [1]. These procedures are critical for cell type annotation, principally through the identification of marker genes concentrated in specific clusters. An alternative approach for cell-type annotation, circumventing the clustering process, employs label transfer from a reference dataset for the annotation of individual cells [2].

However, both strategies have intrinsic limitations. Clusters are not inherently indicative of cell types. Although unsupervised clustering can capture signals related to cell type identity, it does not necessarily ensure differentiation, particularly in the case of closely related cell types. This limitation stems from the so-called ‘curse of dimensionality’, inherent noise in scRNA-seq data due to limited mRNA capture, and gene dropouts [3]. Moreover, clusters frequently signify functional similarity rather than explicit cell type identity. Cell populations with similar functional properties or developmental trajectories often appear as continuous transcriptional landscapes rather than discrete clusters, challenging the fundamental assumption that clustering can effectively separate all biologically meaningful cell types. These challenges amplify when attempting to deep phenotype the data, i.e., discern subsets of similar cell types. The alternate strategy, individual cell annotation using methods like SingleR, struggles with the low sensitivity of scRNA-seq data, rendering annotation based on a restricted set of genes.

Given their high-dimensionality, noise, and sparsity, scRNA-seq datasets pose a significant challenge in analysis. Dimensionality reduction, through deconvolving gene expression into discrete latent spaces, often aids in obtaining a manageable low-dimensional data representation. Principal component analysis (PCA) is the most common approach for dimensionality reduction in scRNA-seq analysis and serves as the default method in popular analysis packages. While PCA effectively captures global variance structure, it has several limitations for scRNA-seq data: it assumes linear relationships between features, is sensitive to outliers, and importantly, cannot leverage prior biological knowledge about cell types. Furthermore, batch effects often dominate the variance in scRNA-seq data, causing PCA to emphasize technical rather than biological variation. Integration methods like Harmony [4] attempt to address batch effects by aligning similar cell populations across datasets. While effective for straightforward cases, these methods often struggle with complex scenarios involving multiple batches and closely related cell types. Deep learning approaches such as scVI [5] have shown promise but require extensive computational resources and may lack interpretability.

Non-negative matrix factorization (NMF) has emerged as a powerful alternative. Unlike PCA, the non-negativity constraint and its ability to learn parts-based representations make it particularly well-suited for gene expression analysis [6]. The NMF basis vectors can reveal interpretable patterns such as co-expressed gene modules or cell-type-specific expression programs that might remain hidden when genes are analyzed individually. Several methodologies using NMF for scRNA-seq analysis have emerged, including CASSL [7], which applies base NMF for semi-supervised clustering of partially labeled scRNA-seq datasets, employing recursive k-means clustering until each group contains only a single class of labeled and unlabeled cells; SDCNMF [8], which utilizes an NMF representation to optimize classification while imposing similarity and dissimilarity constraints; and LIGER [9] uses NMF to identify shared and dataset-specific factors that facilitate integration across multiple experiments. Recently, there have been advances in supervised NMF approaches. For example, the DNMF package [10] implements discriminative NMF for improved classification tasks in gene expression data. Other approaches have enhanced performance by incorporating additional prior knowledge [11], such as marker genes, while others have demonstrated the potential of transfer learning by using learned bases from annotated data to factorize new, unannotated datasets [12].

Despite these advances, existing methods still face significant challenges in: (1) effectively balancing noise reduction with preservation of cell type-specific signals, (2) integrating data across batches without compromising biological distinctions, (3) detecting rare cell populations, and (4) maintaining interpretable biological relevance in the reduced dimensions.

Here, we present CellMentor, a novel supervised NMF framework that addresses these challenges by leveraging labeled reference datasets to learn biologically meaningful latent spaces that can be transferred to new datasets. CellMentor introduces innovative approaches for rank selection along with a unique loss function that simultaneously preserves cell type identity and reduces technical variation. Our method leverages labeled reference datasets to learn biologically meaningful latent spaces that can be effectively transferred to new datasets. We demonstrate that CellMentor effectively captures fundamental biological differences between cell types while minimizing technical artifacts, enabling more accurate downstream analyses including cell type identification.

## 2 RESULTS

### 2.1 Overview of CellMentor

The CellMentor framework operates in two phases (Figure 1). In the decomposition phase, CellMentor learns cell type-specific patterns from a labeled reference dataset using supervised NMF. This process yields two matrices: *W* (genes × factors) representing gene combinations or meta-genes, and *H* (factors × cells) containing the cell-specific usage of these factors. For the decomposition phase we developed a supervised framework that incorporates discriminative constraints to preserve cell type identity. The supervised framework incorporates a novel optimization function with key parameters that balance reconstruction accuracy, sparsity, orthogonality, within-class compactness, and between-class separation.

**Figure 1.**
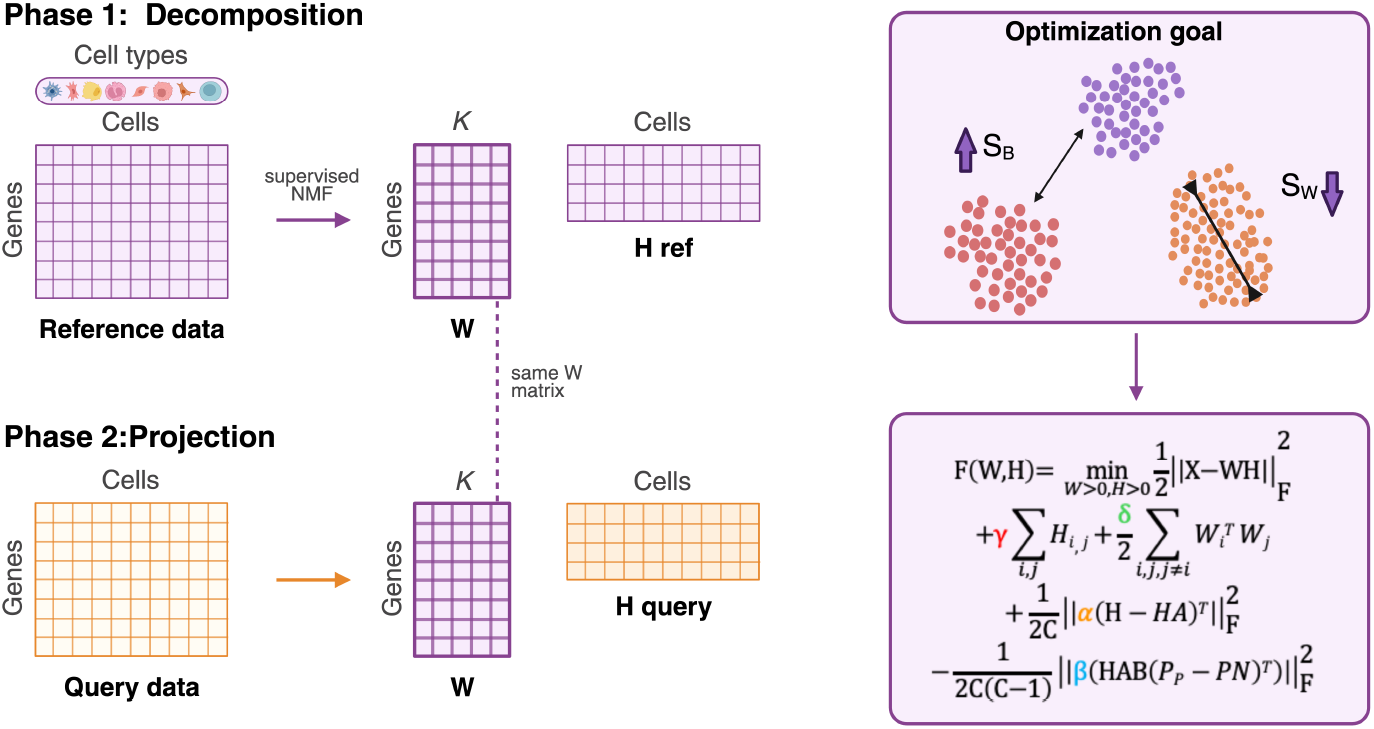
Schematic overview of the CellMentor methodology. The framework operates in two phases: decomposition and projection. In the decomposition phase (top), a labeled reference dataset is factorized using supervised non-negative matrix factorization (NMF) to learn a gene-factor matrix W (genes × K) and a factor-cell matrix H ref (K × cells), where K represents the optimal number of factors determined through biwhitening and eigenvector analysis. The optimization goal (top right) is to simultaneously minimize within-class variation (*S*_*W*_) while maximizing between-class separation (*S*_*B*_). The loss function (bottom right) incorporates four key components: a reconstruction term that ensures accurate representation of the original data, a sparsity term (*γ*) that promotes interpretable factor representations, an orthogonality constraint (*δ*) that minimizes redundancy between factors, a within-class compactness term (*α*) that minimizes variation among cells of the same type, and a between-class separation term (*β*) that maximizes distances between different cell types. In the projection phase (bottom), the learned W matrix is applied to a new query dataset to project it into the same factor space, generating H query that represents the query cells in the biologically meaningful latent space learned from the reference data. This approach enables effective transfer of cell type-specific patterns across datasets while mitigating batch effects.

In the projection phase, CellMentor applies these learned patterns to analyze new datasets. Specifically, we use the *W* matrix learned during decomposition to project query data into the learned latent space. This transfer learning approach enables robust cell type identification across experimental batches without requiring retraining, making it particularly valuable for integrating new datasets with established reference atlases. The resulting reduced-dimensional representation effectively manages technical variation, facilitating downstream analyses such as cell type identification and characterization of rare populations.

### 2.2 Development and Optimization of the Decomposition Phase

A critical challenge in dimensionality reduction is determining the optimal number of dimensions (factors, K) that capture true biological signal while excluding technical noise. In the context of NMF, this rank determines the number of meta-genes (columns) in the *W* matrix and their corresponding usage patterns (rows) in the *H* matrix. CellMentor addresses this through a novel approach combining the biwhitening method with eigenvector analysis [13]. The biwhitening method transforms the data matrix such that both its rows and columns have unit variance, enabling more accurate estimation of the matrix rank. Using controlled simulations with known ground truth, we demonstrated that standard approaches often overestimate the required dimensions by conflating technical variation with biological signal (Supplementary Figure 1).

Our approach extends the biwhitening framework by incorporating two key components. First, we evaluate different scaling techniques (Poisson and quadratic variance) before applying biwhitening, selecting the approach that minimizes the distance between the empirical eigenvalue distribution and the theoretical Marchenko-Pastur (MP) distribution. The quadratic variance scaling technique consistently showed superior performance in our simulations. Second, we refine the rank estimation through eigenvector localization analysis. Using simulated data where ground truth is known, we showed that delocalized eigenvectors often correspond to technical variation rather than biological signal. By setting a significance level of 0.01 for eigenvector localization, we effectively filtered out dimensions dominated by technical noise while preserving biologically meaningful variation.

The core innovation of CellMentor lies in its supervised learning framework, which guides dimensionality reduction to explicitly preserve cell type identity. Our approach incorporates discriminative constraints into the NMF optimization, designed to maximize separation between different cell types while maintaining coherence within cell populations. This is achieved through a novel optimization function that balances four key components: (1) a reconstruction term that ensures the factorization accurately represents the original data, (2) a within-class compactness term that minimizes variation among cells of the same type, with adaptive weights that increase for cells farther from their cluster centroid, (3) a between-class separation term that maximizes distances between different cell types, with stronger weights for similar cell types that are more likely to be confused, and (4) a sparsity constraint that promotes interpretable factor representations. Each component is controlled by tunable parameters that adapt to local and global data characteristics.

### 2.3 Evaluation of CellMentor in Simulated Data

We first evaluated CellMentor using a systematic simulation framework designed to test its performance under controlled conditions with increasing levels of complexity. Using Splatter [14], we generated two sets of simulated scRNA-seq datasets with known ground truth cell types and batch effects.

In our first simulation experiment, we generated 10 distinct scRNA-seq datasets of increasing difficulty, each containing two batches and six cell types with varying degrees of technical noise (Methods). For each simulation, we designated one batch as reference data for the decomposition phase and one as query data for the projection phase, mimicking real-world scenarios where reference and query data come from different experimental conditions. We compared CellMentor against a comprehensive set of established dimensionality reduction methods: Principal Component Analysis (PCA, the standard approach used in Seurat [15]), several NMF and PCA variants (ICA [16], NMF [17], pCMF [18], GLM-PCA [19]), and scANVI [5].

The reduced dimensional representations obtained from different methods showed varying degrees of success in preserving cell type structure (Figure 2A and Supplementary Figure 2). In the most challenging simulation (simulation #10), standard PCA-based dimensionality reduction resulted in suboptimal cell type separation, with multiple cell types merged into single clusters (using default parameters). Particularly visible is a central cluster containing multiple cell populations, making it difficult to accurately identify distinct cell types without manual adjustment of clustering parameters. Similarly, scANVI created continuous structures but with imprecise boundaries between cell types, which also led to suboptimal clustering results. In contrast, CellMentor’s reduced dimensional space showed well-separated cell type clusters with near-perfect correspondence between clusters and true cell type labels, requiring no adjustment of clustering parameters to achieve accurate cell type identification. Across all ten simulations, CellMentor achieved a mean ARI of 0.97 (range: 0.82-1.0), while PCA showed a mean ARI of 0.75 (range: 0.1-1.0) and scANVI reached a mean ARI of 0.48 (range: 0.15-0.69).

**Figure 2.**
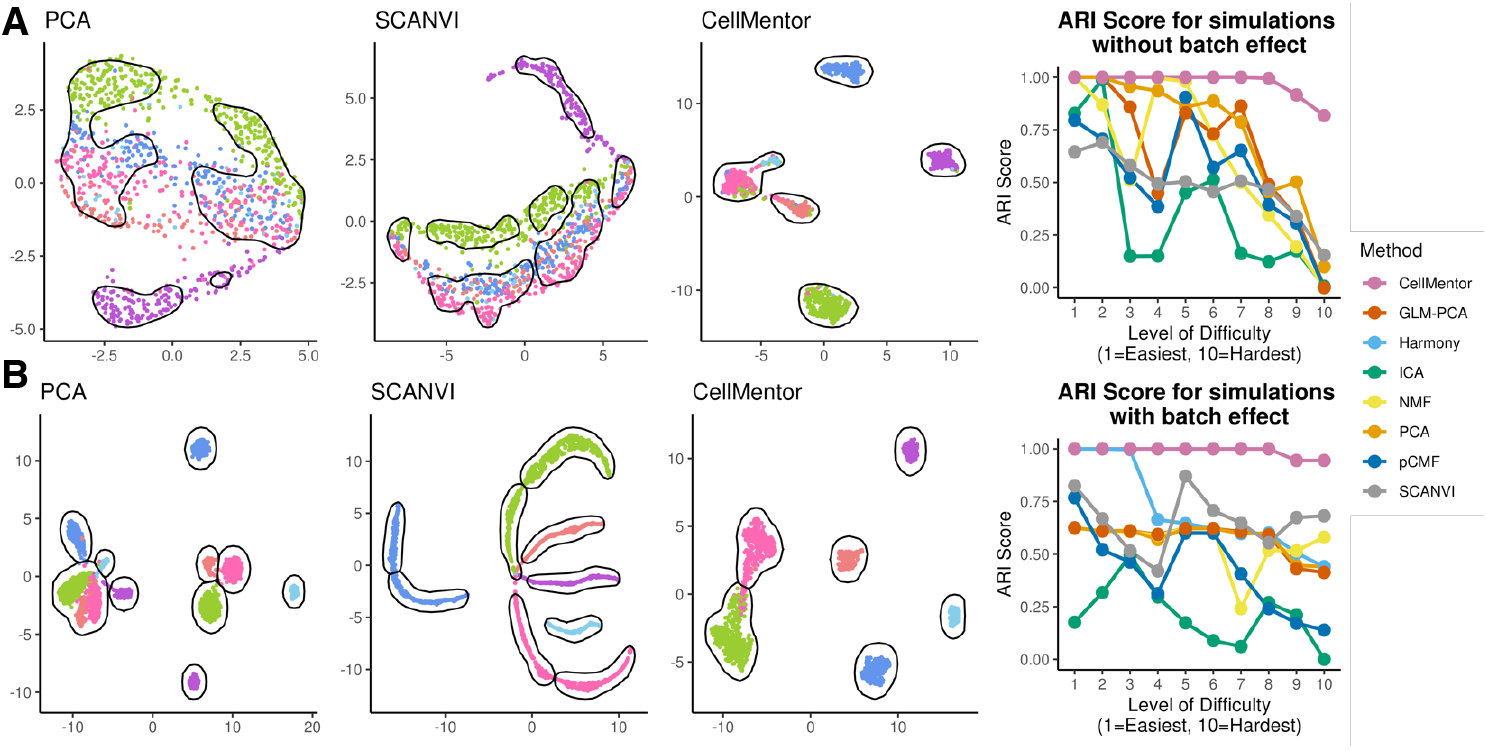
Evaluation of CellMentor using simulated data. **A**. Comparison of dimensionality reduction methods without batch effects. The query data was analyzed using PCA, scANVI and CellMentor and visualized using UMAP. Simulation #10 is presented, all other simulations can be found in Supplementary Figure 2-3. Each reference and query data have 6 cell types. Plots are colored by cell type with identified clusters outlined in black. The graph (right) shows ARI scores across 10 simulations of increasing difficulty. **B**. Similar to A, but for simulations with two batches in the reference and two in the query.

Quantitative assessment using the adjusted rand index (ARI) metric confirmed these observations (Figure 2A). Competing methods showed highly variable performance even in simpler scenarios (simulations 1-5), with several methods displaying dramatic performance drops as early as simulation 3. As simulation complexity increased, all comparison methods continued to deteriorate, showing erratic performance and a general downward trend. By simulation 10, none of the comparison methods achieved an ARI above 0.25, with some dropping to near zero, indicating poor concordance between cluster assignments and true cell types. In stark contrast, CellMentor maintained perfect clustering performance across simulations 1-8, and still achieved substantially higher performance in the most challenging scenarios (simulations 9-10), demonstrating its robustness to increasing levels of technical noise and biological complexity.

In our second simulation experiment, we increased the complexity by generating datasets with four batches instead of two, using two batches for reference and two for query (Figure 2B and Supplementary Figure 3). This scenario represents the common challenge of integrating data across multiple experimental batches while preserving cell type identity. We included Harmony [4], a popular batch correction method, in our comparisons for this more complex scenario. All comparison methods exhibited substantial variability and generally lower performance across simulations. Standard dimensionality reduction methods like PCA showed particularly poor performance in later simulations, with ARI scores dropping below 0.2 and a mean ARI of only 0.57 (range: 0.44-0.63) across all simulations. Even methods specifically designed for batch integration, such as Harmony, failed to maintain consistent performance, with a mean ARI of 0.71 (range: 0.44-1.0) and declining to around 0.4 in the most challenging scenarios. In stark contrast, CellMentor demonstrated remarkably stable and superior performance in this multi-batch scenario, maintaining a mean ARI of 0.99 (range: 0.95-1.0) across all simulations, with values at or near perfect for simulations 1-8 and showing only a slight decrease to approximately 0.95 for the most challenging simulations (9-10).

To further evaluate CellMentor’s capabilities and limitations, we conducted additional simulation experiments exploring challenging scenarios in single-cell analysis (Figure 3A). First, we tested its behavior when encountering novel cell types by excluding one cell type from the reference dataset while maintaining it in the query data (Figure 3B). While CellMentor did not create a completely distinct cluster for this novel cell type (which would be expected for a supervised approach with no prior knowledge of this class), it still showed some ability to separate these cells from known cell types in the query data, demonstrating partial robustness to incomplete reference data.

**Figure 3.**
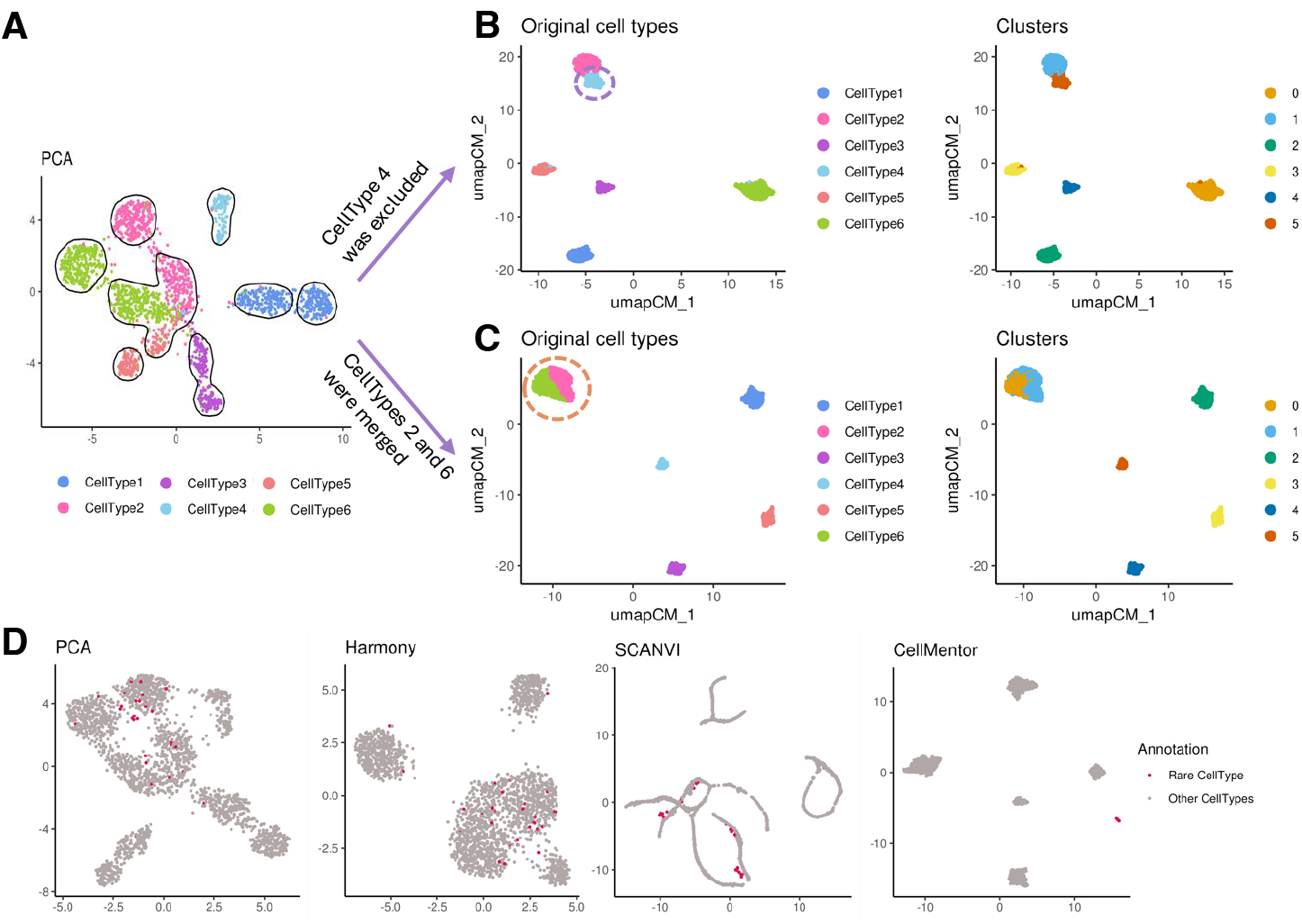
Application of CellMentor to challenging scenarios. **A**. PCA-based UMAP visualization of the simulation dataset with all six cell types, showing how different cell types are naturally distributed. **B**. Performance on novel cell type detection: cell type 4 was removed from the reference data but retained in the query. UMAP visualization of the query data showing original cell types (left) and clusters (right). The purple dashed circle highlights where cell type 4 appears in the embedding despite not being present in the reference. **C**. Performance with incomplete reference annotation: cell types 2 and 6 were merged in the reference data. UMAP visualization showing original cell types (left) and clusters (right). The orange dashed circle shows where cell types 2 and 6 appear together but maintain some separation despite the merged annotation. **D**. Detection of rare cell populations: UMAP visualizations of query data from different dimensionality reduction methods, with rare cells highlighted in red. PCA, Harmony and scANVI scatter rare cells across the embedding, while CellMentor successfully clusters them into a distinct population.

In a related experiment, we tested how CellMentor would handle cases where reference data annotation is not sufficiently granular - a common challenge in single-cell analysis (Figure 3C). We relabeled cells of one type as another type in the reference dataset, effectively merging two distinct populations under a single label. Despite training with this “corrupted” reference data, CellMentor still managed to partially separate these cell populations in the query data, suggesting that the method can detect underlying biological variation even when reference annotations do not fully capture it.

Finally, we assessed the ability to handle rare cell populations by simulating a scenario where one of the cell types represented only 1% of cells across all batches (Figure 3D). In this challenging setting, other tested dimensionality reduction methods failed to identify the rare population, with these cells appearing scattered throughout the embedding space with no distinct clustering. Standard approaches like PCA and even integration methods like Harmony were unable to preserve the signal from this rare population. CellMentor, however, leveraging its supervised framework and the presence of this rare population in the reference data, successfully maintained separation of this rare cell type in the query data, forming a distinct, coherent cluster.

### 2.4 Application to Diverse Tissue Types

We next evaluated CellMentor on real data using three pairs of datasets from different tissue types. Each pair consisted of a reference dataset for training and an independent query dataset for validation: pancreatic tissue (Baron [20]/Muraro [21]), peripheral blood mononuclear cells (PBMCs, [22]/Ding [23]), and melanoma samples (Tirosh [24]/Jerby-Arnon [25]). After standard preprocessing (Methods), we applied CellMentor and competing methods to each tissue type.

For pancreatic tissue analysis (Figure 4B), we observed that PCA, scANVI, and CellMentor, performed well in separating the major pancreatic cell types. This relatively high performance across these methods likely reflects the distinct transcriptional profiles of different pancreatic cell types, making this a more straightforward classification problem. During our analysis, we identified 16 cells that were annotated as alpha cells in the original Muraro dataset but were clustered with the beta cells according to CellMentor. We further investigated these cells using SingleR for independent annotation, which confirmed they expressed gene signatures more consistent with beta cells than alpha cells. This finding demonstrates how our approach can help identify potential misannotations in published datasets.

**Figure 4.**
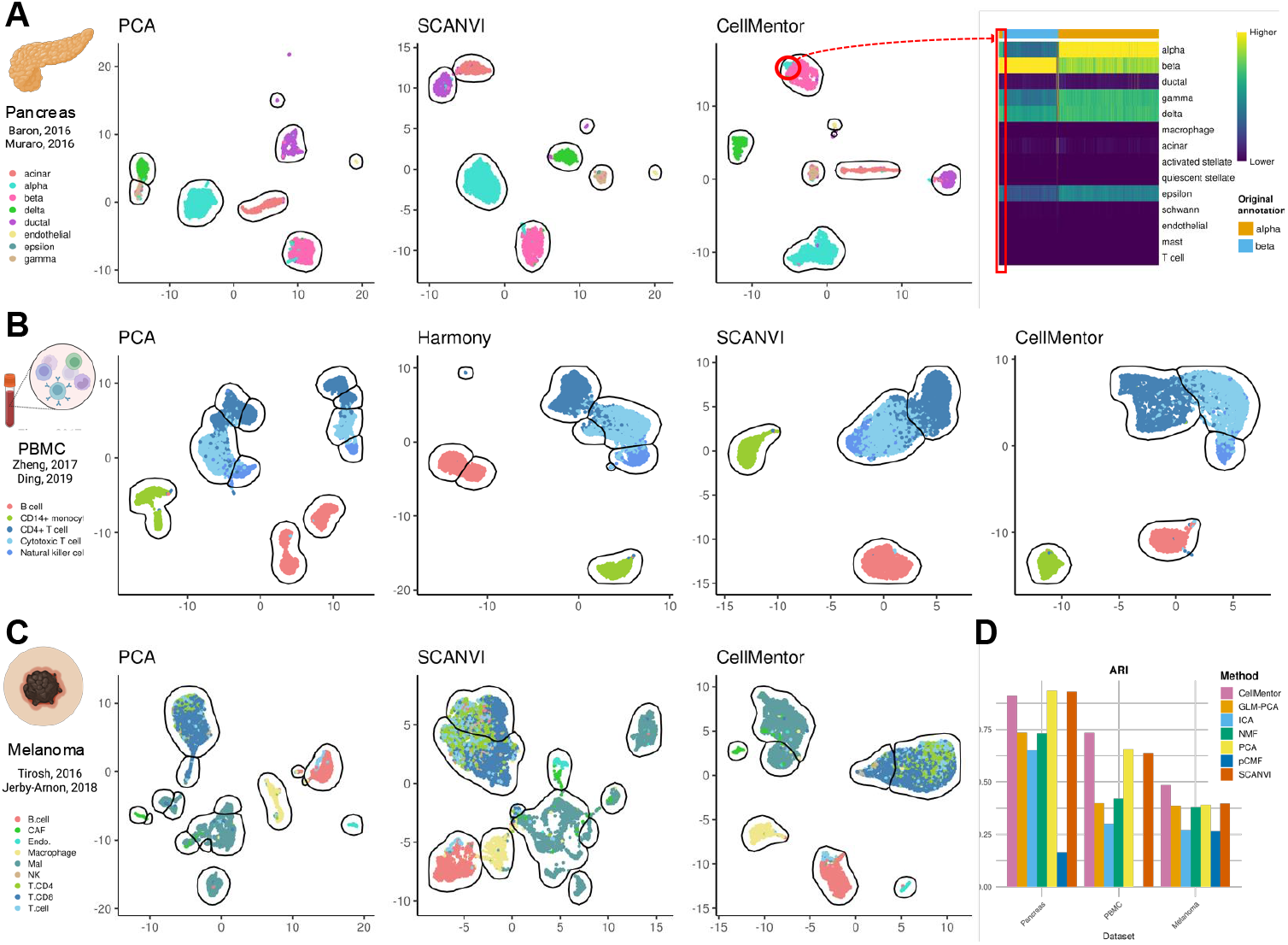
Evaluation of CellMentor on real datasets. **A**. UMAP visualization of PBMC data from Ding using Zheng as the reference. PCA-based UMAP shows distinct clustering by patient batch rather than cell type. Harmony and scANVI integrate batches but with limitations in cell type separation. CellMentor effectively integrates batches while maintaining clear cell type boundaries. **B**. UMAP visualization of pancreatic tissue data from Muraro using Baron as the reference. The red circle and SingleR scores heatmap highlight cells annotated as alpha cells in the original study that cluster with beta cells. **C**. UMAP visualization of melanoma data using Tirosh as reference and Jerby-Arnon as query. CellMentor shows superior separation of tumor and immune cell populations compared to PCA and scANVI. **D**. Quantitative evaluation using Adjusted Rand Index (ARI) across the three tissue types. CellMentor shows strong performance across all datasets, with the highest ARI scores in pancreas and melanoma tissues, while performing comparably to the best alternative methods in the PBMC dataset.

In PBMC analysis, the query dataset (Ding) contained samples from two different individuals, presenting a common challenge in single-cell analysis where batch effects often dominate technical variation (Figure 4A). The PCA-based dimensionality reduction failed to integrate these batches, creating distinct clusters for each patient rather than grouping cells by biological type. While both Harmony and scANVI successfully integrated the batches, they showed limitations in preserving cell type structures - Harmony created multiple clusters for B cells and CD8+ T cells, while scANVI failed to clearly isolate natural killer (NK) cells. In contrast, CellMentor not only successfully integrated the batches but also generated well-defined clusters that corresponded more precisely to known cell types. Notably, none of the methods, including CellMentor, created completely distinct clusters for CD4+ T cells, CD8+ T cells, and NK cells. This likely reflects the biological reality that these cell types exist on a transcriptional continuum rather than as discrete populations, highlighting a limitation of discrete clustering approaches for certain cell types.

The melanoma datasets (Figure 4C) presented the most challenging scenario, with complex mixtures of tumor and infiltrating immune cells that typically do not separate as cleanly as cells from normal tissues. This scenario represents many real-world tumor datasets where cellular states exist along continuums rather than as discrete types. In this context, CellMentor demonstrated superior performance compared to both PCA and scANVI, creating more distinct and biologically meaningful clusters. While PCA showed poor separation between similar cell types and scANVI created overlapping clusters for several cell populations, CellMentor maintained clearer boundaries between different cell types while still effectively integrating cells from different patients.

Quantitative evaluation using the ARI confirmed CellMentor’s strong performance across all three tissue types (Figure 4D). CellMentor achieved the highest ARI scores in both pancreas and melanoma datasets. For the PBMC dataset, CellMentor performed comparably to the best alternative methods, with scANVI and PCA showing similar ARI scores. These results demonstrate CellMentor’s effectiveness across diverse tissue types and experimental settings, highlighting its consistent performance across different scenarios while showing particular strength in datasets with distinct transcriptional profiles like the pancreatic tissue.

## 3 DISCUSSION

Single-cell RNA sequencing (scRNA-seq) is a powerful tool for investigating cellular heterogeneity, enabling analysis of gene expression at the single-cell level. The substantial expansion of high-throughput biological assays, such as scRNA-seq, has led to the generation of massive volumes of data. The ability to analyze individual cells from diverse tissues has allowed for comprehensive and precise comparisons, but it has also intensified the complexity and scalability requirements of data analysis. Dimension reduction is one of the most essential preprocessing steps in scRNA-seq data analysis, aimed at reducing data complexity and eliminating noise. Nevertheless, the extraction of biologically significant molecular features across samples remains a significant challenge in this process.

Several conventional dimension reduction techniques, such as PCA, primarily emphasize technical features that contribute to the greatest variation in the data, often highlighting batch effects rather than biological signals, as demonstrated in our analysis of multi-batch simulations and PBMC data. In contrast, our NMF-based method with appropriate regularization encourages the discovery of informative and discriminative factors that effectively represent the samples in a condensed manner while preserving the inherent differences between classes.

In this study, we introduced CellMentor, a novel approach for dimensionality reduction, clustering, and classification tasks in single-cell transcriptomics. Our systematic evaluation using both simulated and real data demonstrated that CellMentor outperforms existing methods in several key aspects. First, CellMentor showed robustness to increasing levels of technical noise and biological complexity, maintaining near-perfect clustering performance in simulations where other methods degraded significantly. Second, it demonstrated superior ability to handle batch effects while preserving biological signals, a critical capability for integrating data across multiple experiments. Third, CellMentor excelled at detecting rare cell populations, successfully identifying populations representing just 1% of cells where other methods failed completely.

When applied to diverse real-world datasets from pancreas, PBMC, and melanoma tissues, CellMentor demonstrated consistent advantages over existing approaches. In particular, its performance in the challenging melanoma dataset, where tumor heterogeneity creates complex transcriptional landscapes, highlights its value for analyzing clinically relevant samples. Moreover, the case of potentially misannotated cells in the pancreatic dataset demonstrates how improved dimensionality reduction can reveal insights that might be missed with traditional approaches.

A key innovation in CellMentor is its supervised framework that explicitly preserves cell type identity while managing technical variation. Unlike methods that rely solely on variance-based approaches, CellMentor incorporates biological knowledge through its discriminative loss function, ensuring that the reduced dimensions capture fundamental biological differences between cell types. Furthermore, the non-negativity constraint and parts-based representation of NMF provide biological interpretability that is often lacking in other approaches, with meta-genes frequently corresponding to meaningful gene expression programs.

While CellMentor demonstrates significant advantages, several limitations should be considered for future development. The factorization approach employed in CellMentor relies on reference labels to extract latent patterns. The discriminative regulatory terms impose the differentiation of various cell types on the inferred basis vectors. Moreover, the hyperparameter update is designed to assign higher weights to distant cells and linked but distinct groups during the minimization task, promoting compact and distinguishable classes. These characteristics make the algorithm highly sensitive to the accuracy of the reference annotation labels. Incorrect cell labeling can result in the identification of misleading factors that unnecessarily separate cells of the same type or merge different groups. Furthermore, since the learned factors and corresponding constant vectors are used for the projection and annotation of target datasets, these confounding factors impact the accuracy of the analysis for unlabeled data.

To mitigate the bias resulting from mislabeling, one approach is identifying outliers and assigning them lower weights in the optimization problem. This approach aims to reduce the influence of mislabeled cells or groups. However, it is important to note that excluding cells that deviate from common patterns may hinder the identification and classification of rare subcellular types. Therefore, further investigation is required to explore and address this issue in future studies.

As single-cell atlases continue to grow and reference-based analysis becomes increasingly common, approaches like CellMentor that effectively transfer knowledge across datasets while preserving biological signals will play an increasingly important role in extracting meaningful insights from complex single-cell data. The combination of biological interpretability, technical robustness, and supervised learning framework addresses key challenges in the field, making CellMentor a valuable addition to the single-cell analysis toolkit.

## 4 METHODS

### 4.1 CellMentor Algorithm Overview

CellMentor is a supervised dimensionality reduction framework for single-cell RNA sequencing data that operates in two phases: decomposition and projection. The method leverages labeled reference datasets to learn biologically meaningful reduced dimensions that can be effectively transferred to new datasets. In the decomposition phase, given a labeled reference dataset *X*_*re f*_, CellMentor employs supervised non-negative matrix factorization to decompose the data into two matrices:

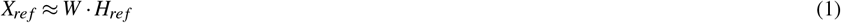

where *W* (genes × factors) represents combinations of genes (meta-genes) and *H*_*re f*_ (factors × cells) contains the cell-specific usage of these factors. The decomposition is guided by cell type labels through a supervised learning framework that incorporates discriminative constraints to preserve cell type identity.

In the projection phase, CellMentor uses the learned *W* matrix to analyze new query datasets. For a query dataset *X*_*query*_, we solve a non-negative least squares problem using the Lawson-Hanson algorithm:

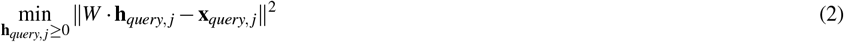

where **x**_*query, j*_ is the *j*^*th*^ column of *X*_*query*_, and **h**_*query, j*_ is the *j*^*th*^ column of *H*_*query*_. The resulting *H*_*query*_ matrix represents the query data in the same reduced dimensional space as the reference data.

The decomposition phase comprises three critical components: (1) determining the optimal number of factors through a biwhitening approach combined with eigenvector analysis, (2) establishing robust matrix initialization strategies, and (3) implementing a supervised framework that incorporates discriminative constraints. These components work together to ensure the learned dimensions capture true biological variation while minimizing technical noise.

### 4.2 Supervised NMF Framework

The core of CellMentor is a supervised NMF framework that incorporates both discriminative constraints and pattern expression requirements. Our objective function combines four key components:

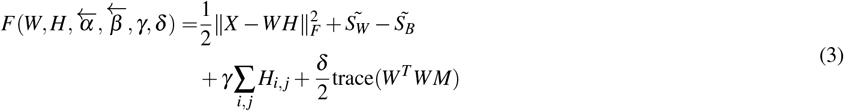

The first term represents the reconstruction error. The second and third terms, 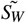 and 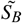, are within-class and between-class scatter matrices:

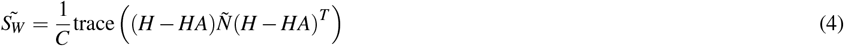

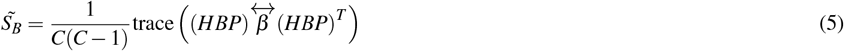

where *C* is the number of cell types, *A* is a block diagonal matrix encoding class membership, and *P* encodes class pair combinations. The fourth and fifth terms enforce sparsity and orthogonality constraints respectively.

The parameters 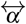 and 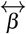 are diagonal matrices that adaptively weight the influence of individual cells and class pairs:

- 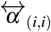 increases for cells farther from their class centroid
- 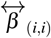 increases for class pairs that are frequently confused

The optimization is solved through multiplicative update rules:

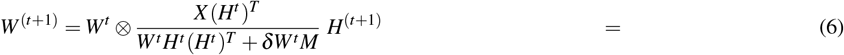

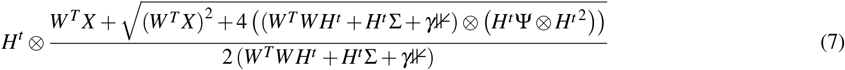

Parameters are tuned using a train-validation split approach. For *γ* and *δ*, we select from predefined ranges based on clustering accuracy on the validation set. For 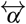 and 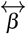, we employ an iterative approach that updates weights based on classification errors and cell-centroid distances.

### 4.3 Key Components of the Decomposition Phase

#### 4.3.1 Rank Selection

CellMentor employs a two-step approach to determine the optimal number of factors (rank). First, we use the biwhitening method [13] to transform the data matrix such that both rows and columns have unit variance. We evaluate different scaling techniques (Poisson and quadratic variance) before biwhitening, selecting the approach that minimizes the distance between the empirical eigenvalue distribution and the theoretical Marchenko-Pastur (MP) distribution.

Second, we refine the rank estimation through eigenvector localization analysis. We evaluate each eigenvector with an eigenvalue exceeding the MP distribution using the two-sided Kolmogorov-Smirnov test, testing whether its components follow a Gaussian distribution with zero mean and variance 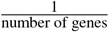. The final rank is determined by the number of localized eigenvectors (p-value ¡ 0.01), effectively filtering out dimensions dominated by technical noise while preserving biological variation.

#### 4.3.2 Matrix Initialization

Due to the non-convex nature of NMF, initialization of W and H matrices is crucial. We implemented five initialization strategies:

Random-based approaches:

- Uniform random initialization in the interval [*ε*, max(X)]
- Up/down-regulated metagene initialization combining random initialization with classification prior knowledge

Structured approaches:

- Non-negative double singular value decomposition (NNDSVD) [26]
- Gene clustering-based initialization using fuzzy c-means
- Cell clustering-based initialization using k-means

After comprehensive evaluation across multiple datasets, we found that the up/down-regulated metagene approach consistently produced the most robust results. This approach has been set as the default initialization method in CellMentor. However, all initialization strategies remain implemented in the package, allowing users to select alternative methods if desired for specific applications.

#### 4.3.3 Parameter Optimization

The supervised framework requires careful tuning of four key parameters 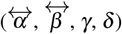. We employ a hierarchical approach:

1. Global parameters (*γ, δ*) are selected from predefined ranges using grid search and validation set performance
2. Cell-specific weights 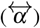 are computed based on distance to class centroid and class prediction errors
3. Class-pair weights 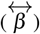 are updated based on confusion frequencies between cell types

Parameters are normalized to maintain balance between within-class and between-class terms, with user-defined constants controlling their relative influence.

### 4.4 Simulation Framework

We used Splatter (v1.26.0) [14] to generate synthetic scRNA-seq datasets for systematic evaluation of CellMentor. Each simulation consisted of either two or four batches (1,000 cells per batch) and six cell types with varying degrees of technical noise. Ten distinct simulation scenarios were created with progressively increasing difficulty to thoroughly test algorithm robustness. For each simulation, one or two batches were designated as reference data for the decomposition phase and the remaining batches as query data for the projection phase, mimicking real-world scenarios where reference and query data come from different experimental conditions. The simulations were designed with systematically increasing difficulty (simulations 1-10) by manipulating key parameters. Early simulations (1-2) featured more balanced cell distributions, minimal batch effects (batch.facLoc = 0.05, batch.facScale = 0.1), low biological variation (bcv.common = 0.2), and uniform dropout rates. Middle simulations (3-6) introduced stronger batch effects, higher biological variation (bcv.common increasing to 0.5), and heterogeneous dropout patterns across batches. The most challenging simulations (7-10) incorporated high biological variation (bcv.common = 0.7-0.8), stronger batch effects (batch.facScale up to 0.5), higher outlier probability (out.prob = 0.1), more heterogeneous dropout patterns, and highly skewed cell type proportions with up to 30% of cells in one type and as low as 3-7% in rare cell types. All simulation parameters are detailed in Supplementary Table 1.

To test specific challenges, we designed additional simulation experiments. In one set of simulations, we tested rare cell type detection by setting one cell type to represent 1% of the total population across all batches. In another set, we evaluated novel cell type identification by excluding one cell type from the reference data while maintaining it in the query data. Performance was evaluated using two clustering metrics: Adjusted Rand Index (ARI) and Normalized Mutual Information (NMI), comparing cluster assignments in the reduced dimensional space to true cell type labels.

### 4.5 Real Data Analysis

We evaluated CellMentor on three pairs of published scRNA-seq datasets spanning different tissue types, each consisting of a reference dataset for training and an independent query dataset for validation:

- Pancreatic tissue: Baron human pancreas dataset (reference, n=8,569 cells) and Muraro pancreas dataset (query, n=2,042 cells). For feature selection, we supplemented variable genes with known pancreatic markers including GCG, INS, SST, PPY, GHRL, KRT19, CPA1, and PDGFRB. In the Muraro dataset, mesenchymal cells were excluded to maintain consistency with other pancreatic datasets.
- Peripheral blood mononuclear cells (PBMCs): Zheng dataset (reference, n=6,932 cells) and PBMC Single Cell Atlas dataset (query, n=7,120 cells). Cell type-specific markers including CD19, CD79A, MS4A1, CD3D, CD8A, IL7R, and NKG7 were included in the feature list. In the PBMC dataset, we excluded cells with high mitochondrial gene expression (¿5%) and abnormally high feature counts (¿4,000) as indicators of poor quality.
- Melanoma: Tirosh melanoma dataset (reference, n=4,097 cells) and Jerby-Arnon dataset (query, n=6,879 cells), with feature selection supplemented by known markers including CD2, CD3D, CD19, CD79A, and melanoma-specific genes such as MIA and TYR.

Data preprocessing included log-transformed normalization 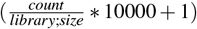, feature selection using the SingleR (v2.4.1) [2] trainSingleR algorithm, and scaling using root-mean-square-deviation (RMSD). For each tissue type, we retained only genes shared between reference and query datasets (pancreas: 15,290 genes; PBMC: 15,752 genes; melanoma: 22,667), selecting the most variable genes for analysis (pancreas: 755; PBMC: 487; melanoma: 562).

All analyses were performed on distinct single cells, with each cell representing an independent measurement. No repeated measurements of the same cells were conducted across any of the analyses.

Data analysis was performed using R (v4.3.2). CellMentor utilizes the following packages: methods (v4.3.2), Matrix (v1.6-5), sparsesvd (v0.2-2), RMTstat (v0.3.1), SingleR (v2.4.1), MLmetrics (v1.1.3), nnls (v1.5), parallel (v4.3.2), progress (v1.2.3), cluster (v2.1.4), skmeans (v0.2-17), irlba (v2.3.5.1), lsa (v0.73.3), data.table (v1.16.2), ggplot2 (v3.5.1), magrittr (v2.0.3), tibble (v3.2.1), Seurat (v5.1.0), utils (v4.3.2), matrixStats (v1.4.2).

### 4.6 Comparison with Existing Tools

We systematically compared CellMentor against six state-of-the-art dimensionality reduction methods across identical simulation datasets and real-world pairs of reference-query data. All methods were implemented with consistent parameters (20 latent dimensions) and evaluated using standardized workflows. Principal Component Analysis (PCA) was implemented using Seurat (v5.1.0) [15] with 2,000 variable features. Independent Component Analysis (ICA) was performed using the fastICA package (v1.2-5.1) [16] with tolerance set to 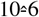 and the logcosh contrast function. Non-negative Matrix Factorization (NMF) was executed using the NNLM package (v0.4.4) with Lee’s multiplicative update algorithm, 500 maximum iterations, and 10^−6^ convergence tolerance. Probabilistic Count Matrix Factorization (pCMF) [18] was implemented with both zero-inflation and sparsity parameters enabled using pCMF package (v1.2.1). Generalized Linear Model PCA (GLM-PCA) was applied directly to count data using the glmpca package (v0.2.0) [19] with Poisson family modeling. For single-cell variational inference (scVI), we utilized the Python-based scvi-tools (v1.2.1) [5] framework with three hidden layers (256, 128, and 64 nodes), batch correction, and 250 training epochs with 10^−3^ learning rate. All dimension reduction outputs were visualized through uniform manifold approximation and projection (UMAP) using identical parameters (`n neighbors=15`, `min dist=0.3`) via the Seurat RunUMAP function. For both simulated and real data, we assessed each method’s performance using a standardized workflow: (1) application of dimensionality reduction, (2) UMAP visualization of latent space, (3) graph-based clustering with multiple resolution parameters (0.1-0.5), and (4) evaluation of clustering quality against ground truth labels (simulations) or trusted annotations (real data). Clustering performance was quantified through Adjusted Rand Index (ARI) and Normalized Mutual Information (NMI) using the aricode package (v1.0.3).

## ACKNOWLEDGMENTS

This work was funded by the Israel Science Foundation (1543/21) and Azrieli Faculty Fellowship.

## Author contributions

D.A. conceived the study, supervised all analyses, and wrote the manuscript. O.H. developed the mathematical framework and core algorithm, implemented the initial codebase, and performed preliminary benchmarking analyses. E.P. conducted simulation studies, analyzed real datasets, transformed the code into a software package, performed additional benchmarking, and generated the figures. All authors reviewed and approved the final manuscript.

## Data Availability

All data used in this study are publicly available. The PBMC datasets are available from Zheng et al. (2017) through the 10X Genomics website (https://support.10xgenomics.com/single-cell-gene-expression/datasets) and from Ding et al. (2020) through GEO accession number GSE132044. The pancreatic tissue datasets from Baron et al. (2016) and Muraro et al. (2016) are available through GEO accession numbers GSE84133 and GSE85241, respectively. The melanoma datasets from Tirosh et al. (2016) and Jerby-Arnon et al. (2018) are available through GEO accession numbers GSE72056 and GSE115978. Simulated datasets were generated using the Splatter R package.

## Code Availability

The CellMentor package developed in this study is freely available as an open-source R package in a public GitHub repository (https://github.com/petrenkokate/CellMentor). Complete documentation, including installation instructions and usage examples, is provided in the repository. All analysis code, benchmark tests, and scripts for reproducing the figures presented in this paper are available in a separate GitHub repository (https://github.com/petrenkokate/CellMentor_paper).

## REFERENCES

1. Heumos, L. et al. Best practices for single-cell analysis across modalities. en. Nat. Rev. Genet. 24, 550–572 (Aug. 2023).

2. Aran, D. et al. Reference-based analysis of lung single-cell sequencing reveals a transitional profibrotic macrophage. en. Nat. Immunol. 20, 163–172 (Feb. 2019).

3. Qiu, P. Embracing the dropouts in single-cell RNA-seq analysis. en. Nat. Commun. 11, 1169 (Mar. 2020).

4. Korsunsky, I. et al. Fast, sensitive and accurate integration of single-cell data with Harmony. en. Nat. Methods 16, 1289–1296 (Dec. 2019).

5. Gayoso, A. et al. A Python library for probabilistic analysis of single-cell omics data. en. Nat. Biotechnol. 40, 163–166 (Feb. 2022).

6. Esposito, F. A Review on Initialization Methods for Nonnegative Matrix Factorization: Towards Omics Data Experiments. Mathematics 9, 1006. ISSN: 2227-7390. https://www.mdpi.com/2227-7390/9/9/1006 (Apr. 2021).

7. Seal, D. B., Das, V. & De, R. K. CASSL: A cell-type annotation method for single cell transcriptomics data using semi-supervised learning. Applied Intelligence 53, 1287–1305. ISSN: 0924-669X. 10.1007/s10489-022-03440-4%20 https://link.springer.com/10.1007/s10489-022-03440-4 (Jan. 2023).

8. Zhu, Y. L., Yuan, S. S. & Liu, J. X. Similarity and Dissimilarity Regularized Nonnegative Matrix Factorization for Single-Cell RNA-seq Analysis. Interdisciplinary Sciences – Computational Life Sciences 14, 45–54. ISSN: 18671462. https://link.springer.com/article/10.1007/s12539-021-00457-0 (Mar. 2022).

9. Welch, J. D. et al. Single-Cell Multi-omic Integration Compares and Contrasts Features of Brain Cell Identity. Cell 177, 1873–1887. ISSN: 00928674. https://linkinghub.elsevier.com/retrieve/pii/S0092867419305045 (June 2019).

10. Jia, Z. et al. Gene Ranking of RNA-Seq Data via Discriminant Non-Negative Matrix Factorization. PLOS ONE 10 (ed Jordan, I. K.) e0137782. ISSN: 1932-6203. https://dx.plos.org/10.1371/journal.pone.0137782 (Sept. 2015).

11. Wu, P. et al. A robust semi-supervised NMF model for single cell RNA-seq data. PeerJ 8, e10091. ISSN: 2167-8359. https://peerj.com/articles/10091 (Oct. 2020).

12. Stein-O’Brien, G. L. et al. Decomposing Cell Identity for Transfer Learning across Cellular Measurements, Platforms, Tissues, and Species. Cell Systems 8, 395–411. ISSN: 24054712. 10.1016/j.cels.2019.04.004%20 https://linkinghub.elsevier.com/retrieve/pii/S2405471219301462 (May 2019).

13. Landa, B., Zhang, T. T. C. K. & Kluger, Y. Biwhitening Reveals the Rank of a Count Matrix. SIAM Journal on Mathematics of Data Science 4, 1420–1446. ISSN: 2577-0187. http://arxiv.org/abs/2103.13840%20 10.1137/21M1456807 (Dec. 2022).

14. Zappia, L., Phipson, B. & Oshlack, A. Splatter: simulation of single-cell RNA sequencing data. en. Genome Biol. 18 (Dec. 2017).

15. Hao, Y. et al. Dictionary learning for integrative, multimodal and scalable single-cell analysis. en. Nat. Biotechnol. 42, 293–304 (Feb. 2024).

16. Hyvärinen, A. & Oja, E. Independent component analysis: algorithms and applications. en. Neural Netw. 13, 411–430 (May 2000).

17. Gaujoux, R. & Seoighe, C. A flexible R package for nonnegative matrix factorization. en. BMC Bioinformatics 11, 367 (July 2010).

18. Durif, G., Modolo, L., Mold, J. E., Lambert-Lacroix, S. & Picard, F. Probabilistic count matrix factorization for single cell expression data analysis. en. Bioinformatics 35, 4011–4019 (Oct. 2019).

19. Townes, F. W., Hicks, S. C., Aryee, M. J. & Irizarry, R. A. Feature selection and dimension reduction for single-cell RNA-Seq based on a multinomial model. en. Genome Biol. 20, 295 (Dec. 2019).

20. Baron, M. et al. A Single-Cell Transcriptomic Map of the Human and Mouse Pancreas Reveals Inter-and Intra-cell Population Structure. Cell systems 3, 346–360. ISSN: 2405-4712. https://pubmed.ncbi.nlm.nih.gov/27667365/. (Oct 2016).

21. Muraro, M. J. et al. A Single-Cell Transcriptome Atlas of the Human Pancreas. Cell Systems 3, 385–394. ISSN: 24054712. https://linkinghub.elsevier.com/retrieve/pii/S2405471216302927 (Oct. 2016).

22. Zheng, G. X. Y. et al. Massively parallel digital transcriptional profiling of single cells. Nature Communications 8, 14049. ISSN: 2041-1723. https://www.nature.com/articles/ncomms14049 (Jan. 2017).

23. Ding, J. et al. Systematic comparison of single-cell and single-nucleus RNA-sequencing methods. Nature Biotechnology 38, 737–746. ISSN: 15461696. http://dx.doi.org/10.1038/s41587-020-0465-8 (2020).

24. Tirosh, I. et al. Dissecting the multicellular ecosystem of metastatic melanoma by single-cell RNA-seq. en. Science 352, 189–196 (Apr. 2016).

25. Jerby-Arnon, L. et al. A cancer cell program promotes T cell exclusion and resistance to checkpoint blockade. en. Cell 175, 984–997.e24 (Nov. 2018).

26. Boutsidis, C. & Gallopoulos, E. SVD based initialization: A head start for nonnegative matrix factorization. Pattern Recognition 41, 1350–1362. ISSN: 00313203. https://linkinghub.elsevier.com/retrieve/pii/S0031320307004359 (Apr. 2008).

