## Supplementary Figure 1 for "CellMentor: Cell-Type Aware Dimensionality Reduction for Single-cell RNA-Sequencing Data"

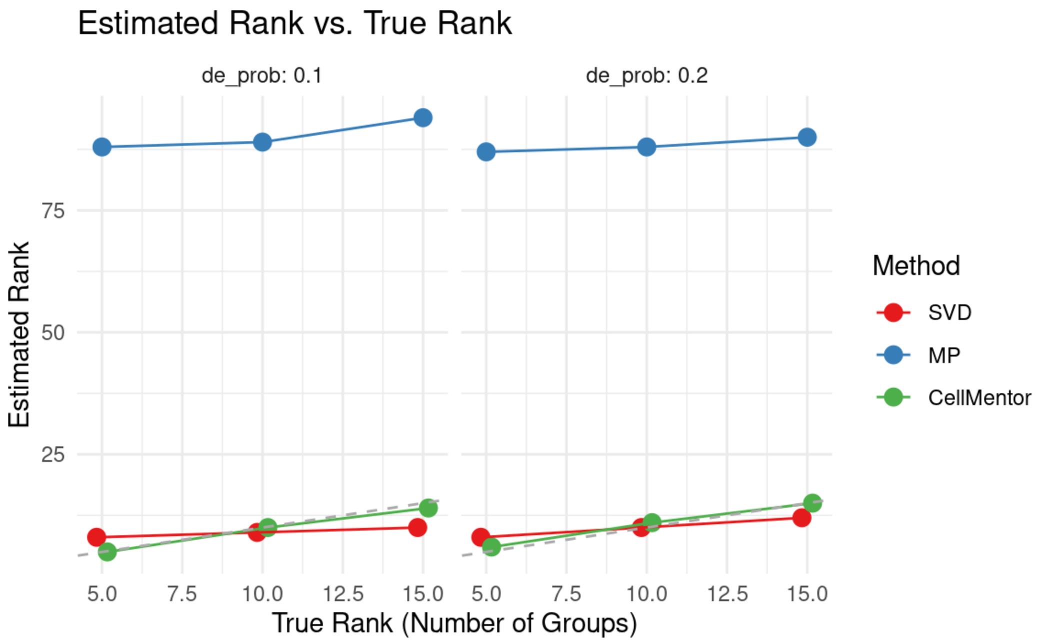


Supplementary figure 1. Comparison of rank estimation methods across varying biological complexity in single-cell RNA-seq data. Estimated rank versus true underlying dimensionality (number of cell groups) for three dimensionality estimation approaches: SVD (red), MP law (blue), and CellMentor (green). The grey line represents the ground truth where estimated rank equals true rank. Simulations were performed using Splatter with differential expression probabilities of 0.1 (left) and 0.2 (right). The MP method substantially overestimates rank in all scenarios (y-axis truncated at 100), while both SVD and CellMentor produce estimates closer to ground truth. CellMentor demonstrates superior performance at higher complexities, particularly with stronger differential expression signals (de_prob: 0.2), highlighting its effectiveness in distinguishing biological variation from technical noise in complex transcriptomic landscapes.
