## Supplementary Figure 2 for "CellMentor: Cell-Type Aware Dimensionality Reduction for Single-cell RNA-Sequencing Data"

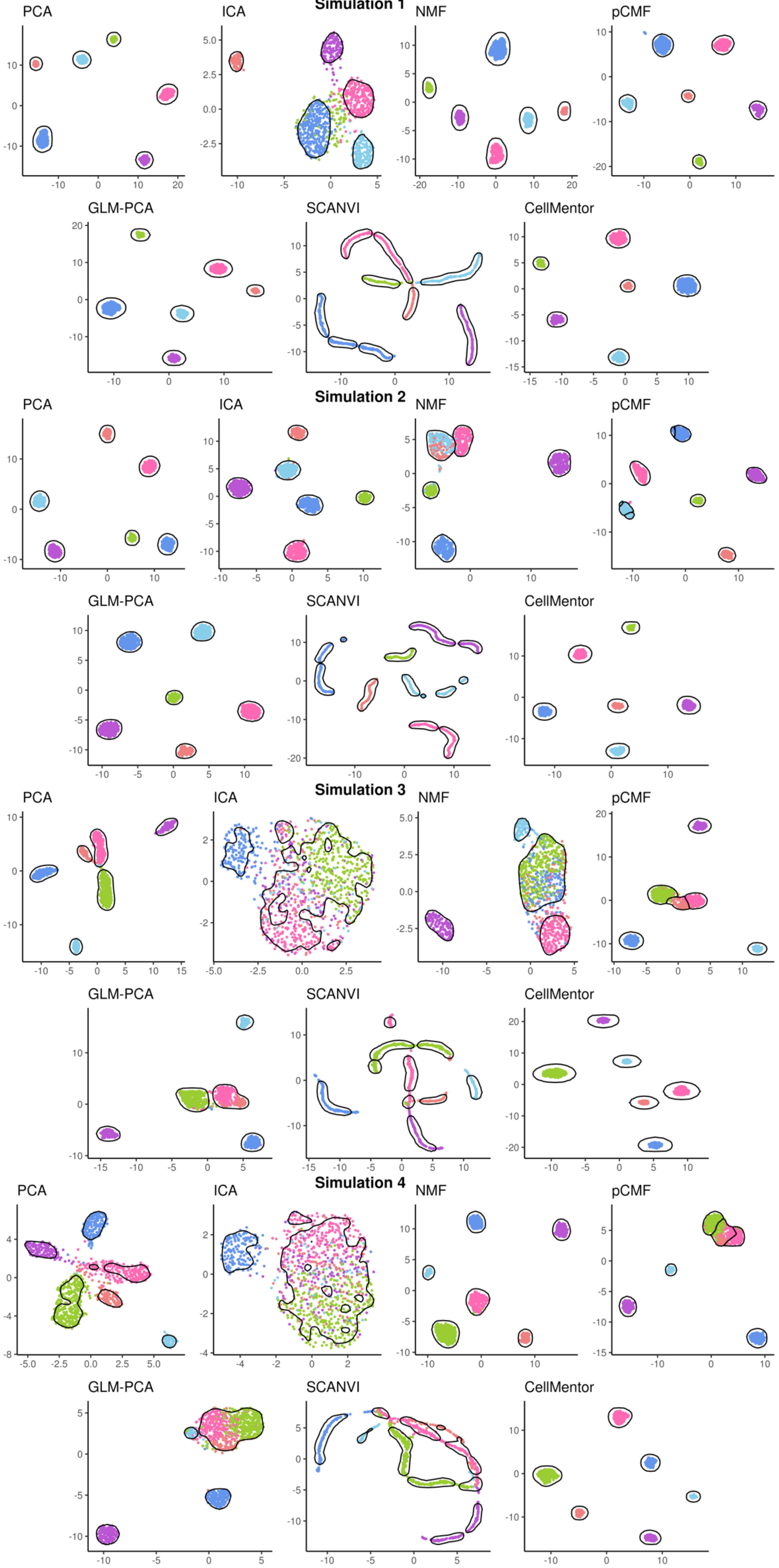

Simulation 1
PCA
ICA
NMF
pCMF
GLM-PCA
SCANVI
CellMentor
Simulation 2
PCA
ICA
NMF
pCMF
GLM-PCA
SCANVI
CellMentor
Simulation 3
PCA
ICA
NMF
pCMF
GLM-PCA
SCANVI
CellMentor
Simulation 4
PCA
ICA
NMF
pCMF
GLM-PCA
SCANVI
CellMentor

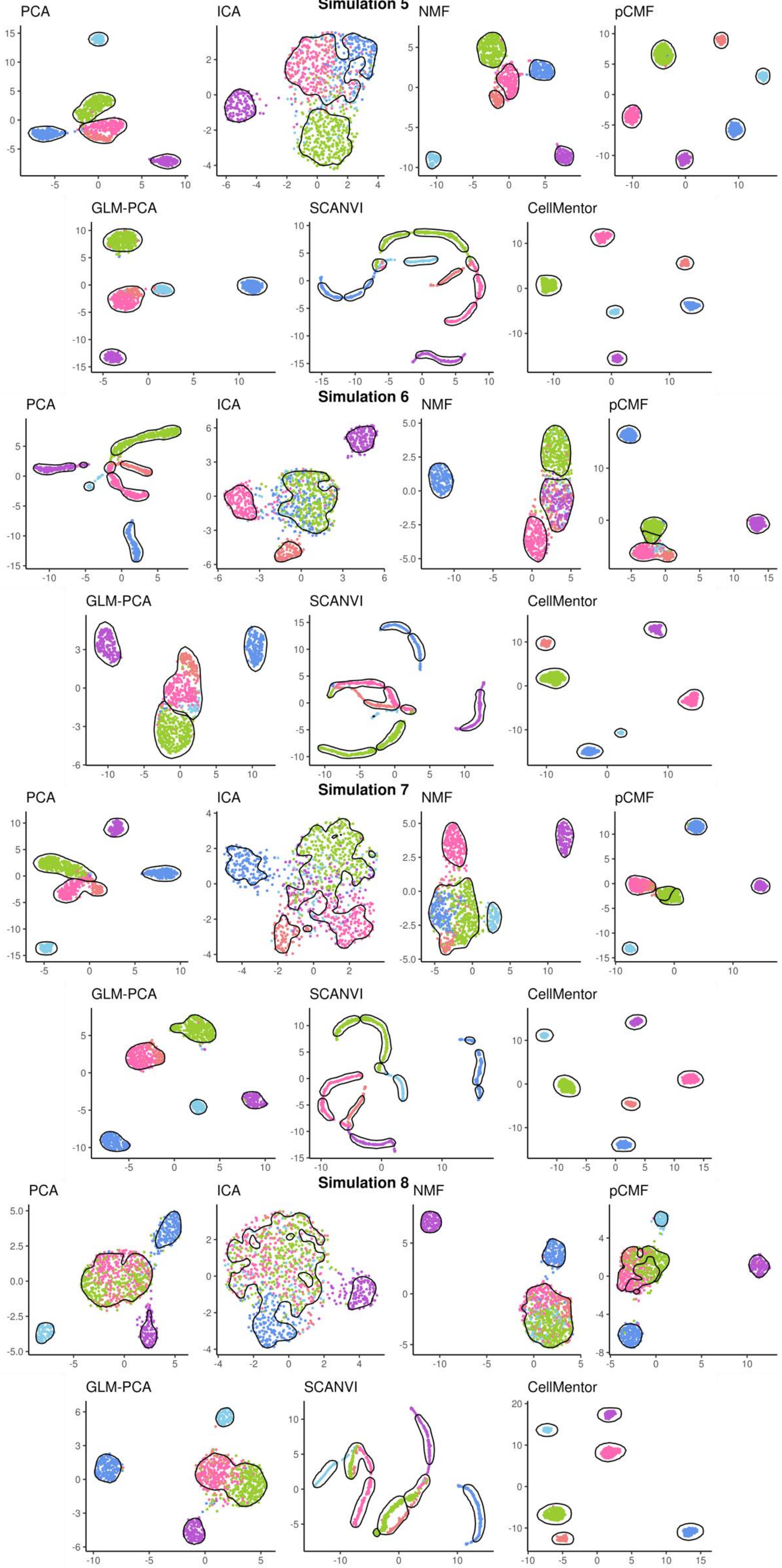
Simulation 5
PCA
ICA
NMF
pCMF
GLM-PCA
SCANVI
CellMentor
Simulation 6
PCA
ICA
NMF
pCMF
GLM-PCA
SCANVI
CellMentor
Simulation 7
PCA
ICA
NMF
pCMF
GLM-PCA
SCANVI
CellMentor
Simulation 8
PCA
ICA
NMF
pCMF
GLM-PCA
SCANVI
CellMentor

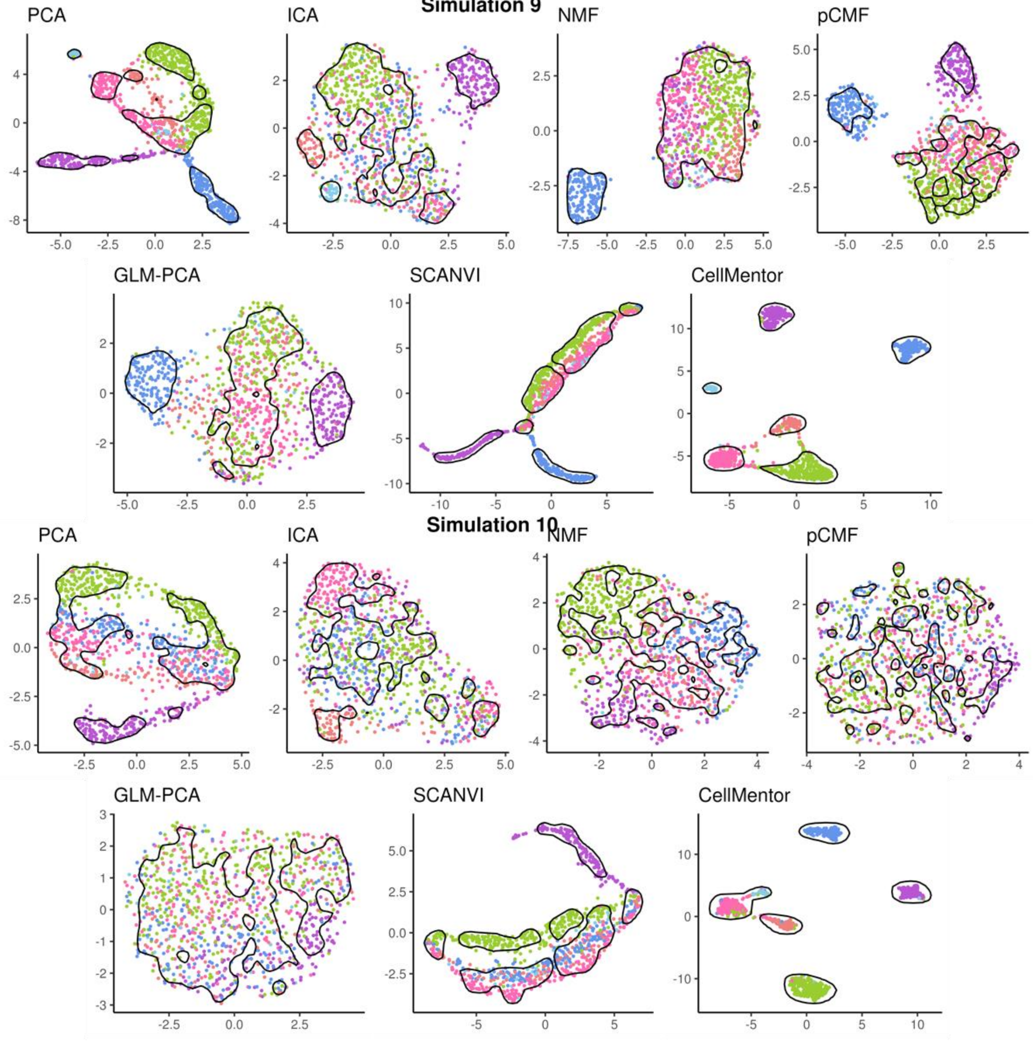


**Supplementary Figure 2: Comparison of dimensionality reduction methods across all simulations without batch effects**

This figure extends the analysis presented in Figure 1A, displaying UMAP visualizations for all 10 simulations with increasing difficulty (simulations 1-10). For each simulation, we compare seven dimensionality reduction and integration methods: PCA, ICA, NMF, pCMF, GLM-PCA, SCANVI, and CellMentor. Each panel shows the projection of cells in two-dimensional space, with colors representing different cell types and black outlines indicating identified clusters.

The visualizations demonstrate how different methods perform in preserving cell type structure as simulation complexity increases. In simpler simulations (lower numbers), most methods successfully separate cell populations. However, as difficulty increases, many conventional methods show deteriorating performance with overlapping clusters and poor cell type separation. CellMentor consistently maintains distinct cell type clusters across all difficulty levels, corresponding to its higher ARI scores as shown in Figure 1A. SCANVI also shows characteristic curved manifold structures but with varying separation quality across simulations. This comprehensive comparison highlights the robust performance of CellMentor in maintaining cell type identity information even under challenging conditions.
