## Supplementary Figure 3 for "CellMentor: Cell-Type Aware Dimensionality Reduction for Single-cell RNA-Sequencing Data"

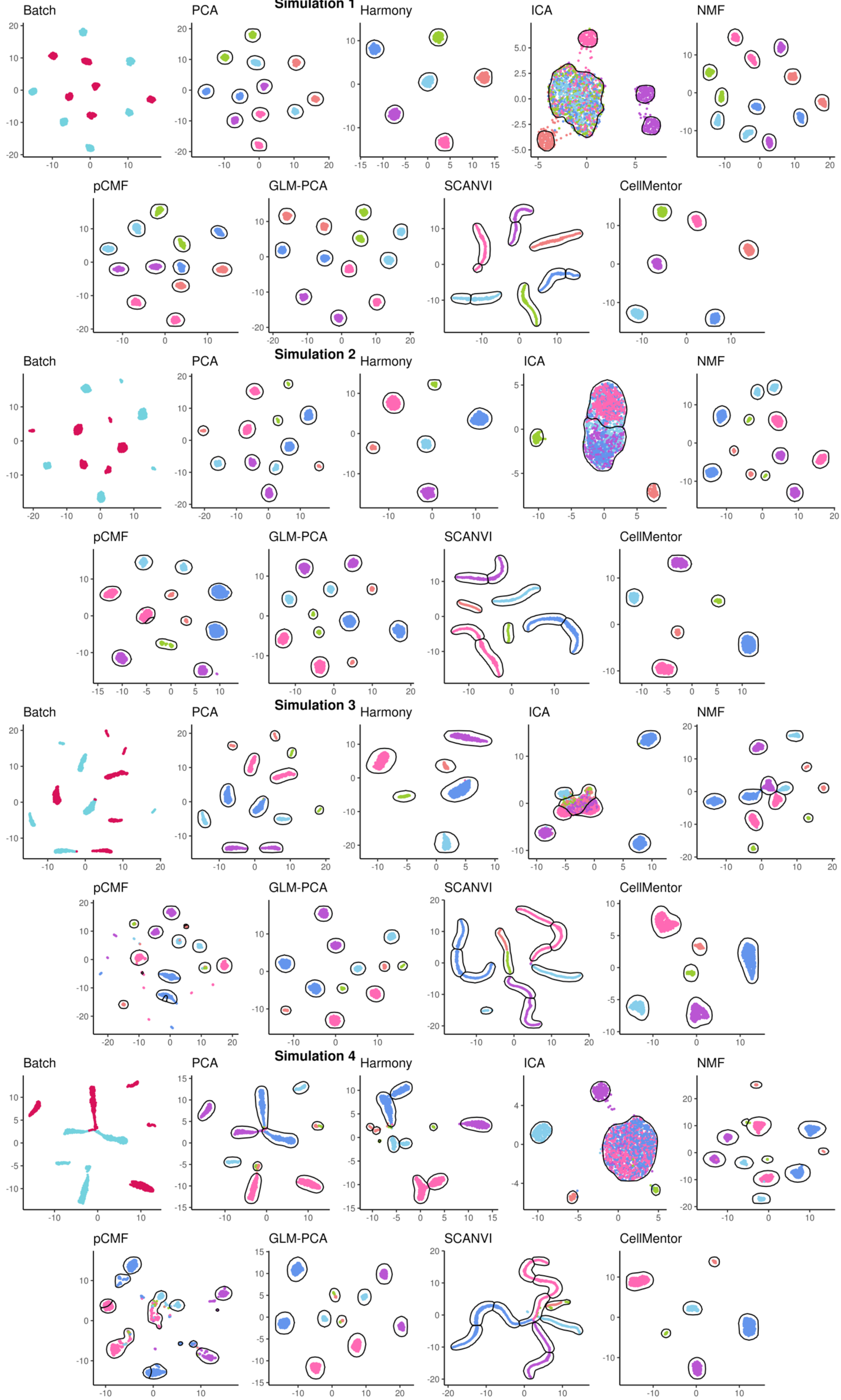

Simulation 1
Batch
PCA
Harmony
ICA
NMF
pCMF
GLM-PCA
SCANVI
CellMentor
Simulation 2
Batch
PCA
Harmony
ICA
NMF
pCMF
GLM-PCA
SCANVI
CellMentor
Simulation 3
Batch
PCA
Harmony
ICA
NMF
pCMF
GLM-PCA
SCANVI
CellMentor
Simulation 4
Batch
PCA
Harmony
ICA
NMF
pCMF
GLM-PCA
SCANVI
CellMentor

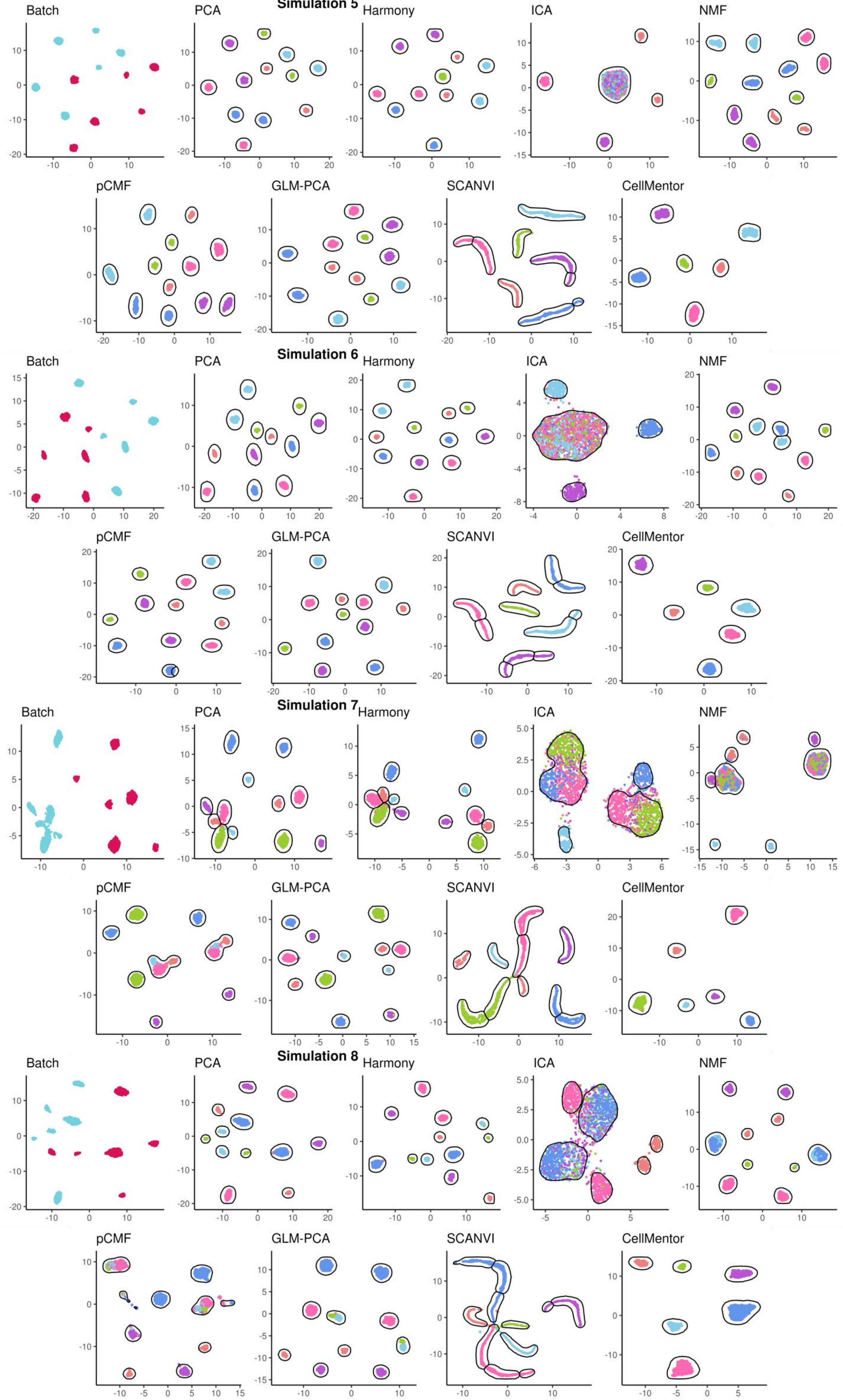

Simulation 5
Batch
PCA
Harmony
ICA
NMF
pCMF
GLM-PCA
SCANVI
CellMentor
Simulation 6
Batch
PCA
Harmony
ICA
NMF
pCMF
GLM-PCA
SCANVI
CellMentor
Simulation 7
Batch
PCA
Harmony
ICA
NMF
pCMF
GLM-PCA
SCANVI
CellMentor
Simulation 8
Batch
PCA
Harmony
ICA
NMF
pCMF
GLM-PCA
SCANVI
CellMentor

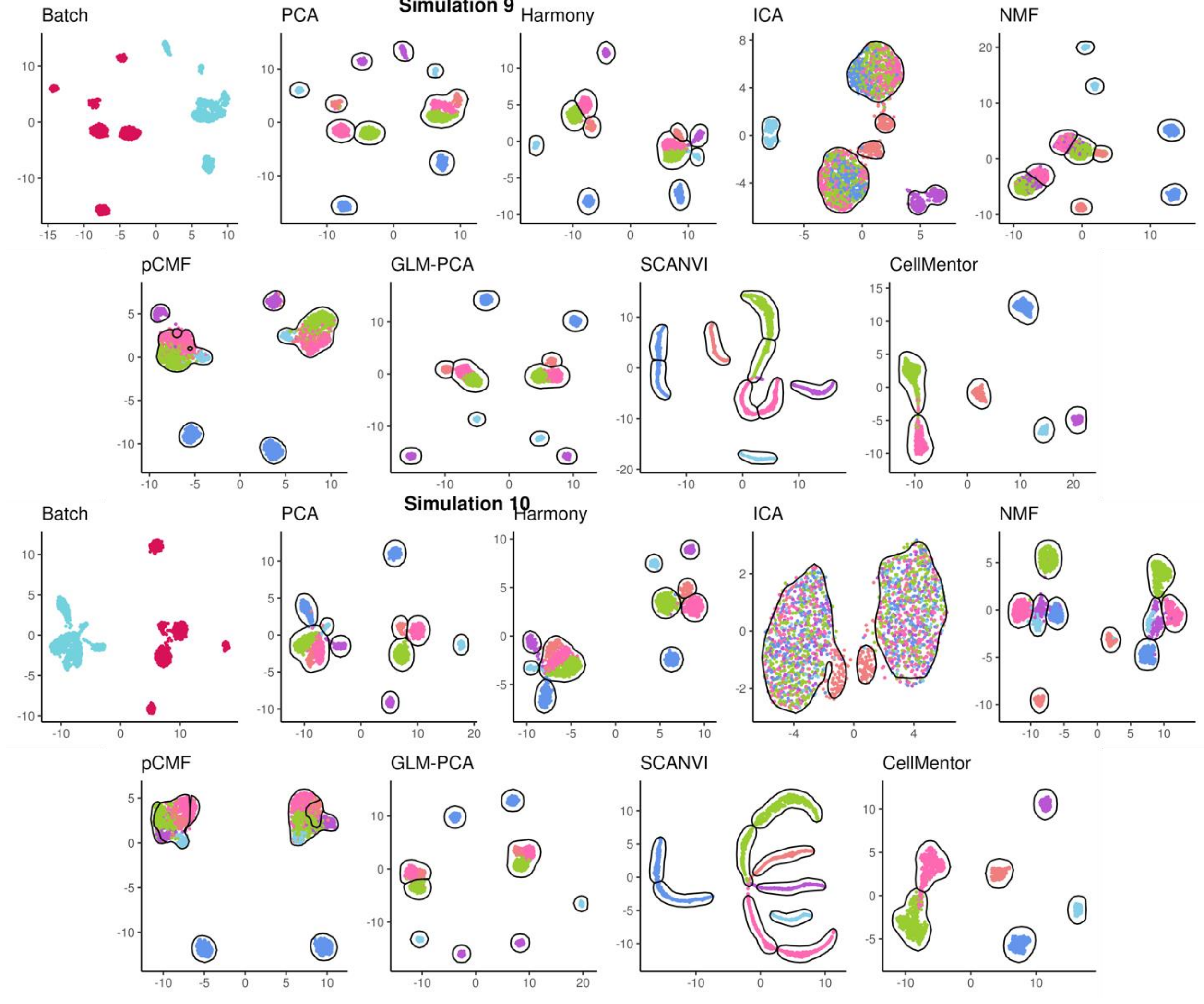


**Supplementary Figure 3: Comparison of dimensionality reduction methods across all simulations with batch effects**

This figure extends Figure 1B, showing UMAP visualizations for all 10 simulations with batch effects of increasing difficulty. Each row presents a simulation with multiple panels: the leftmost "Batch" panel shows data colored by batch (red/blue), while subsequent panels compare eight methods (PCA, Harmony, ICA, NMF, pCMF, GLM-PCA, SCANVI, and CellMentor). Colors represent cell types with identified clusters outlined in black.

As simulation difficulty increases, most methods struggle to simultaneously correct batch effects and preserve cell type identity. Basic methods like PCA and ICA show persistent batch effects, while Harmony improves batch integration but sometimes loses biological signal. CellMentor consistently maintains distinct cell type clusters across all difficulty levels despite batch effects, aligning with its superior ARI scores in Figure 1B. This comparison demonstrates the value of methods that effectively balance batch correction with preservation of biological structure in single-cell data.
